# Catching a wave: on the suitability of traveling-wave solutions in epidemiological modeling

**DOI:** 10.1101/2023.06.23.546298

**Authors:** Anna M. Langmüller, Joachim Hermisson, Courtney C. Murdock, Philipp W. Messer

## Abstract

Ordinary differential equation models such as the classical SIR model are widely used in epidemiology to study and predict infectious disease dynamics. However, these models typically assume that populations are homogeneously mixed, ignoring possible variations in disease prevalence due to spatial heterogeneity. To address this issue, reaction-diffusion models have been proposed as an alternative approach to modeling spatially continuous populations in which individuals move in a diffusive manner. In this study, we explore the conditions under which such spatial structure must be explicitly considered to accurately predict disease spread, and when the assumption of homogeneous mixing remains adequate. In particular, we derive a critical threshold for the diffusion coefficient below which disease transmission dynamics exhibit spatial heterogeneity. We validate our analytical results with individual-based simulations of disease transmission across a two-dimensional continuous landscape. Using this framework, we further explore how key epidemiological parameters such as the probability of disease establishment, its maximum incidence, and its final epidemic size are affected by incorporating spatial structure into SI, SIS, and SIR models. We discuss the implications of our findings for epidemiological modeling and identify design considerations and limitations for spatial simulation models of disease dynamics.

## 1. Introduction

A fundamental understanding of how infectious diseases spread through time and space is crucial for predicting the course of epidemics and guiding potential control measures. Compartmental models have been the workhorse for infectious disease modeling for almost a century [1, 2, 3]. In these models, the population is divided into compartments based on their infection status, such as susceptible (S), infectious (I), and recovered (R) individuals. One of the simplest compartmental models is the so-called SI model, where individuals can only be susceptible or infected [1]. Once infected, an individual remains in that state forever. Two classical extensions of this model are the SIS and SIR models [1]: In the former, infected individuals can return to the susceptible class, while in the latter they can recover, often implying that they are immune to future infections.

Compartmental models are typically studied with ordinary differential equations (ODEs) that describe the expected changes in the proportions of the different population compartments over time [1, 2, 4, 5, 3]. For example, consider a simple SI model where individuals come into contact with each other at a constant rate *r*. Let *i*(*t*) denote the fraction of infectious individuals in the population. Whenever a susceptible individual meets an infected one, it contracts the pathogen with probability *α* (for simplicity, we will use the terms ‘pathogen’ and ‘disease’ interchangeably throughout this paper, recognizing their conceptual distinction). Assuming homogeneous mixing among the individuals, the change in the proportion of infectious individuals over time is given by:

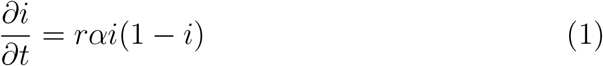

For models with additional compartments, such as a recovered class in the SIR model, the dynamics are described by a set of coupled ODEs. Over the past century, such models have provided invaluable insights into the factors that shape disease transmission by providing conceptual results and threshold quantities (e.g., herd immunity, basic reproduction number, and replacement number) to predict disease spread and the impact of countermeasures, thereby guiding decision-making and public health policy during disease outbreaks [4, 2, 3, 5, 6].

The limitations of ODE models often lie in the underlying assumptions they make about the transmission process. A common assumption of ODE models is homogeneous mixing among individuals [7]. This is rarely the case for real-world populations, which can be structured in many ways [8, 9, 10, 11, 12]. In particular, most populations are geographically structured, so that individuals living close together are more likely to meet than individuals living far apart. Besides spatial structure, kinship and social structure can also have a significant impact on contact patterns [13, 14]. As a result, local disease prevalence can vary substantially between different parts of the population, leading to notable differences in the expected epidemiological dynamics [15, 11, 16]. While classical ODE models have clear advantages for modeling disease transmission, such as their well-understood mathematical properties, computational efficiency, and ability to provide analytical solutions [11], ignoring the influence of population structure in these models can lead to inaccurate conclusions.

In this study, we focus on the potential effects of spatial population structure, which can be incorporated into disease transmission models in several ways. One approach is to use a meta-population model that divides the population into subpopulations or “patches”, each representing a homogeneous population such as a city. The dynamics within the patches, and the coupling between them, can then be described by a system of ODEs [17, 18, 19]. Such coupling could be based on geographic distance, empirical observations such as mobile phone data [20], or geolocated social media posts [21]. Alternatively, individual-or agent-based approaches can be used to explicitly model all individuals in the population and their spatial location [22, 16, 23]. The interactions between individuals (e.g., the rate and strength of contacts) can be regulated by distance-based kernels [5, 24], which can result in highly variable infection probabilities across space [11]. Network models represent another class of approaches that allow for fine-scale modeling of population structure [25]. In these models, each node typically represents an individual, with edges between nodes representing contacts between individuals. A susceptible individual can only contract the disease if an edge connects it to an infectious individual [16, 26]. However, network models are complex, and their analysis may require a large amount of empirical data [16].

For populations inhabiting a continuous landscape, where individuals move primarily within short distances relative to the total geographic range, reaction-diffusion models offer an alternative approach to describe spatial disease dynamics [27, 28]. These models gained popularity in the 1930s, when both Fisher and Kolmogorov used them to explain the spread of a beneficial allele in a one-dimensional habitat [29, 30]. In such models, the beneficial allele can propagate through the population as a “traveling wave”, with a constant and well-defined velocity. Since then, diffusion models have been widely used in mathematical biology to study not only the spread of adaptive alleles but also invasive organisms and infectious diseases [28, 31, 32, 27, 33, 34, 35, 36, 37, 3, 38]. In the context of epidemiology, reaction-diffusion models describe the proportion of infected individuals as a function of space and time. The diffusion term models the random dispersal of infectious individuals or contacts [39], while the reaction term describes the local rates of disease transmission, similar to a compartmental ODE model [27].

In this paper, we use diffusion theory to study under which scenarios a simple compartmental ODE model is sufficient to capture the disease dynamics and when spatial structure needs to be considered. We derive a simple critical threshold *D*_*c*_ for the diffusion coefficient, below which the limited dispersal of individuals significantly affects the disease dynamics. To validate our findings, we perform individual-based simulations using SLiM [40], which allow us to smoothly transition from spatially unstructured to structured populations. We examine the effect of spatial structure on disease dynamics in three classical compartmental transmission models: the SI, SIS, and SIR model [1, 4]. Furthermore, we highlight potential caveats of simulating continuous disease transmission models in a discrete, individual-based, and biologically realistic manner.

## 2. Results

### 2.1. ODE model

Consider a simple Susceptible-Infected (SI) model in a closed population of constant size *N* without birth or death events [1, 4]. There are only susceptible and infectious individuals in the population, and infectious individuals do not recover (i.e., they never leave the infectious compartment). Let *I*(*t*) and *S*(*t*) = *N* − *I*(*t*) denote the total number of infectious and susceptible individuals in the population at time *t*, and *s*(*t*) = *S*(*t*)*/N* and *i*(*t*) = *I*(*t*)*/N* their respective population frequencies with *i*(*t*) + *s*(*t*) = 1. If we assume a homogeneously mixing population in which individuals come into contact with each other at a constant contact rate *r* per time unit, and a disease that establishes in a susceptible individual after contact with an infectious individual with probability *α*, then the change in the proportion of infectious individuals is given by Equation (1). The solution to this ODE model is a logistic growth function [41]:

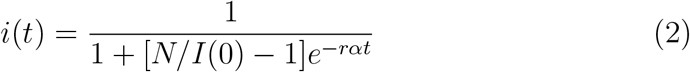

Whenever *I*(0) *>* 0, we have lim_*t*→∞_ *i*(*t*) = 1, meaning that the disease will eventually spread throughout the entire population. Figure 1A provides an illustration of this logistic spread in the ODE model. Note that the parameters *r* and *α* always appear together, with the composite parameter (*rα*)^*−*1^ setting the timescale of the dynamics. However, since *r* and *α* regulate different processes in the context of our individual-based simulation framework described in section 2.4, we do not combine them into one composite variable.

**Figure 1:**
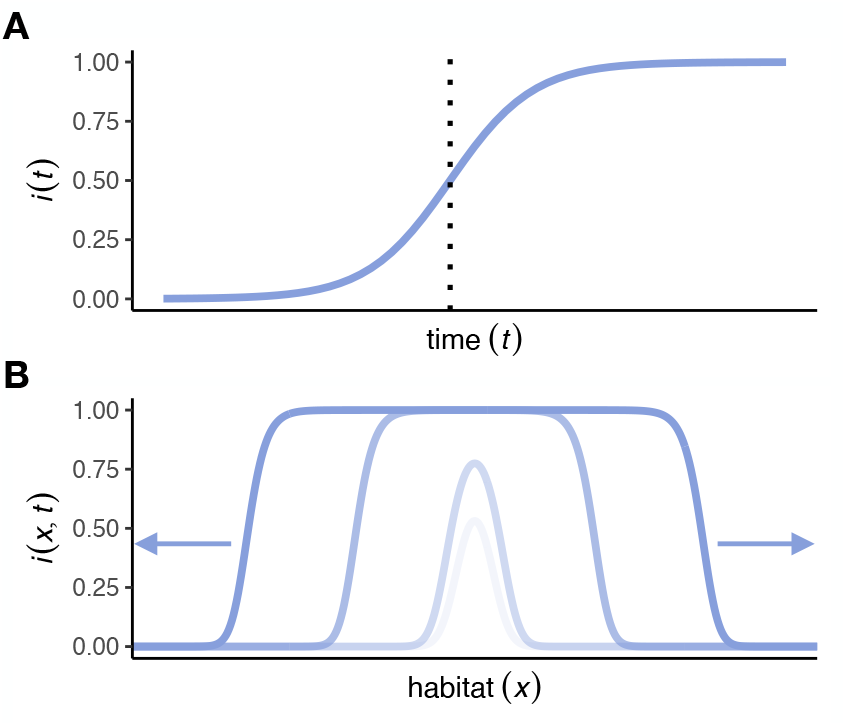
Schematic plots of expected disease spread under the ODE model and a traveling wave solution of the reaction-diffusion model. (A) In the ODE model, the frequency of infected individuals, *i*(*t*), increases logistically according to Equation (2). The vertical dashed lines represent *t*_1*/*2_ as defined in Equation (3). (B) In the diffusion model for a one-dimensional habitat, the disease spreads from an initial release at the center of the habitat via two traveling waves with minimum velocity 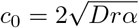 in either direction.

Assuming a single infected individual at time *t* = 0 (*I*(0) = 1) and a sufficiently large population (*N* ≫1), the expected time for the disease to reach 50% frequency in the population is:

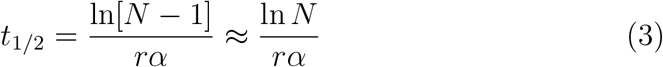

Since *i*(*t*) is S-shaped with the largest growth rate at the inflection point *i*(*t*_1*/*2_) = 1*/*2, the expected time for the disease to spread through the whole population, under the assumptions *N* ≫ 1 and *I*(0) = 1, is therefore:

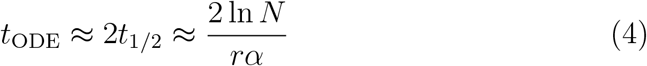

We call *t*_ODE_ the “fixation time” of the ODE model.

### 2.2. Reaction-diffusion model

In contrast to the classical ODE-based compartmental models described above, reaction-diffusion models provide an alternative that explicitly incorporates spatial heterogeneity in disease prevalence [3, 42, 43, 29, 30, 28]. In a diffusion model, the frequency of infected individuals becomes a function not only of time *t* but also of spatial position *x*.

Consider an SI model in which individuals move randomly according to a dispersal kernel that decays with distance at least as quickly as an exponential function [38]. In such a case, we can model dispersal by a diffusion term [38, 29, 44, 28], while disease transmission can be modeled by a reaction term based on the local compartment frequencies. Together, this yields a second-order partial differential equation for the SI diffusion model of the form:

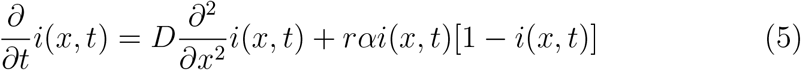

The diffusion coefficient *D* accounts for the random dispersal of individuals. In a one-dimensional habitat, 2*D* = *σ*^2^, where *σ*^2^ is the variance in the expected spatial displacement of individuals per time unit due to their random movement [29, 45]. The reaction term is analogous to the ODE model from Equation (1), except that the frequency of infected individuals now depends on the spatial position *x* as well. Equation 5 is also known as the Fisher-Kolmogorov or Fisher-KPP equation [29, 30, 3].

We can express Equation 5 in dimensionless form by rescaling time, 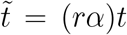, and space, 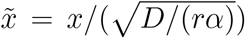 where (*rα*)^*−*1^ is the characteristic timescale of the system:

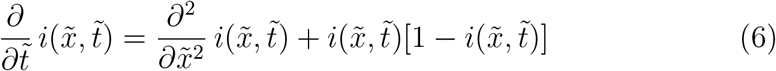

However, in the context of our individual-based simulation model (section 2.4), it is more convenient to maintain the habitat size *L* and the simulation step-size (i.e., “tick”) Δ*t* as the units of space and time. Thus, we retain the parameterization of the Fisher-KPP equation in 5 to facilitate the mapping between the continuous diffusion model and the discrete simulation framework.

Under the assumption that the sum of susceptible and infectious individuals is conserved at a constant carrying capacity, and that there is no change in dispersal due to infection, one solution to Equation 5 is a so-called “traveling wave” [29, 46, 27, 3, 38, 33, 47, 48] in which the disease spreads outward from an initial introduction point. The minimum velocity *c*_0_ of this wave is:

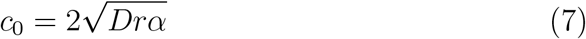

Figure 1B provides an illustration of these dynamics in a one-dimensional habitat, where the disease is introduced at the center and then spreads outward in the form of two symmetric traveling waves to the left and right. The “width” of each wave, i.e., the length of the region between its front (where the frequency of infected individuals is still close to zero) and top (where almost everyone is already infected), is approximately 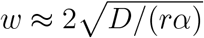 [29].

It is straightforward to extend this diffusion model to two dimensions. Specifically, if dispersal is isotropic, the displacement along the *x* and *y* axes can be treated independently, with each following the one-dimensional model. The diffusion coefficient then equals half the variance of the displacement in each of the *x* and *y* coordinates: 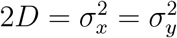. Note that the total displacement in a two-dimensional space, 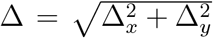, will typically be larger than the displacement in each individual dimension.

In this two-dimensional diffusion model with isotropic dispersal, the disease spreads from an initial introduction point in the form of a growing circle. As the curvature of the wavefront decreases, the velocity at which the radius grows asymptotically approaches the same minimum wave speed *c*_0_ as the traveling wave in the one-dimensional model [8, 49]. If the disease is introduced in a small number of individuals in the center of a square habitat patch with a side length *L* that is much larger than the wave width (which requires 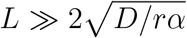 [50]), the wave reaches the corners of the square patch when the circle grows to a radius of 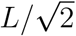.

After an initial phase to establish a wavefront, the expected fixation time of the SI diffusion model is thus governed by the travel time of the wave to reach the corners of the patch, which is approximately:

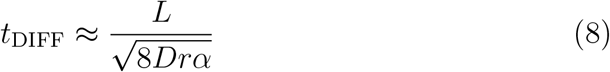

This expression only accounts for the spatial progression and ignores the time it takes for the wavefront to build up and the time for full fixation after the front reaches the habitat boundaries. These are local growth processes. As a simple heuristic, we approximate the additional time needed for these phases by the time for logistic growth of the corresponding ODE model and thus set *t*_fix_ = *t*_DIFF_ + *t*_ODE_ for the total fixation time. More generally, we approximate the time for the disease to reach a given frequency *i*(*t*) in the total population by the sum of the expected time to reach that frequency under a traveling wave (Equation 7) and the time required under logistic growth (Equation 2) alone. We test this heuristic in the simulation below.

### 2.3. Critical threshold

We can now ask under what conditions the spatial component of the disease dynamics cannot be ignored, so that an approximation by the simple ODE model fails. Our exposition of the expected disease fixation time suggests a simple criterion: The spatial component of the dynamics dominates if the travel time of the wave in the diffusion model, *t*_DIFF_, (8), exceeds the time required for homogeneous logistic growth, *t*_ODE_, (3). Setting both equal, we obtain a critical value for the diffusion coefficient:

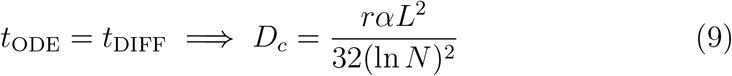

When *D < D*_*c*_ the slow dispersal of individuals through the habitat becomes the limiting factor for the spread of the disease. In this low-dispersal regime, we expect the diffusion model to describe the epidemiological dynamics more accurately than the ODE model, which overestimates the rate of spread.

As *D* becomes of the order of *D*_*c*_, the width of the traveling wave approaches the habitat size, and the disease no longer spreads as a circle with a “sharp” edge [50]. Spatial progression ceases to limit the disease spread, and the fixation time should gradually approach the prediction of the ODE model. In the high-dispersal regime where *D* ≫ *D*_*c*_, diffusion is strong enough that the population is effectively well mixed over the timescale relevant for disease spread, and the ODE model should describe the epidemiological dynamics accurately.

### 2.4. Individual-based simulation framework

To test the theoretical predictions derived above, we implemented an individual-based distributed-infectives simulation model for disease transmission in SLiM (version 4.0) [40]. By varying the dispersal rates of individuals, our simulation model can smoothly transition from spatially unstructured to highly structured populations, allowing us to investigate disease dynamics across the high and low-dispersal regimes, and to study the predicted transition around the critical diffusion coefficient *D*_*c*_. We focused on an abstract, directly transmitted disease in a closed population (i.e., no births or deaths) of *N* = 100, 000 individuals. These individuals move in a continuous two-dimensional spatial domain, modeled as a square habitat patch with a side length of 1 (i.e., *L* = 1) and periodic boundaries to avoid edge effects [51, 15].

The ODE and diffusion models are both continuous time models. Our individual-based simulation model in SLiM, on the other hand, measures time in discrete units of “ticks” (i.e., Δ*t* = 1). Each such “tick” is associated with the opportunity for new infections and recovery of individuals. To minimize differences between the discrete-time simulations and the continuous-time mathematical models, we must ensure that Δ*t* is sufficiently small compared to the characteristic timescale (*rα*)^*−*1^. However, this requirement does not limit the generality of our models. We can freely choose the real-world time interval that a tick in our simulations corresponds to, be it a year or a millisecond. By choosing a sufficiently small period, we can always ensure that Δ*t* ≪ (*rα*)^*−*1^.

The disease is transmitted via local contacts between infectious and susceptible individuals. In each tick, we assume that an individual has contact with all individuals that are currently within its interaction distance δ, which we chose so that each individual has on average 15 contacts per tick (i.e., *r*=15). Note that the average number of contacts per tick is independent of the infection status (i.e., infectious individuals have, on average, as many contacts as susceptible or recovered individuals). For a square habitat with edge length normalized to *L* = 1 and toroidal boundary conditions, and with 100,000 individuals, this corresponds to an interaction radius of δ ≈ 0.00691. Any contact between a susceptible individual and an infectious individual will result in disease transmission with a probability of *α*=0.001, giving (*rα*)^*−*1^ ≈ 66.67 ticks ≫ 1. Once a susceptible individual has contracted the disease, it becomes infectious in the next tick. At the end of each tick, we simulate isotropic dispersal by sampling the *x* and *y* displacements for each individual independently from a normal distribution with mean *µ* = 0 and variance *σ*^2^ = 2*D*. The infection status of an individual has no effect on *D*.

We implemented three compartmental disease transmission models in this simulation framework: (i) the SI model, (ii) the SIS model, and (iii) the SIR model [5]. In the SI model, once an individual is infected, it remains infectious until the end of the simulation. In the SIS model, infected individuals can recover and return to the susceptible class in each tick with probability *γ*. In the SIR model, they recover and gain permanent immunity with probability *γ* in each tick.

Each simulation run is initialized by uniformly distributing 100,000 susceptible individuals throughout the habitat. At tick 25, the disease is introduced into the population by infecting the individual closest to the center of the square habitat (i.e., I(0) = 1). For each simulation, we recorded the number of individuals in each compartment at the beginning of each tick until either the disease spread through the whole population, no infectious individuals were left, or the simulation ran for 50,000 ticks. This upper threshold of 50,000 ticks results in a very mild constraint that is not expected to interfere with the simulated dynamics of SI disease transmission. For our simulation parameters, the disease is expected to fix in the SI model between approximately 1,500–14,000 ticks (Equations 4 and 10).

#### 2.4.1. Low-diffusion limit

A key abstraction of the diffusion approach is that it models a continuous density of individuals throughout the habitat. The relative frequencies of infectious and susceptible individuals are specified at any given location in the habitat, and disease transmission can occur exactly at that point. In our individual-based simulation model, however, a finite number of individuals are distributed across a continuous habitat. It is unlikely that any two of them will ever be at exactly the same point. Thus, one has to decide how close two individuals must be for one to be able to infect the other, which we have implemented by defining an interaction radius δ.

Ideally, δ should be very small (i.e., much smaller than the average dispersal distance of individuals) so that the movement of individuals remains the primary driver of disease spread. Yet, there is a lower limit on δ for any given contact rate in our individual-based simulation model. For example, if we want to ensure that each individual has, say, two contacts on average, this requires some minimum interaction radius so that each individual encounters enough individuals within its interaction radius. In our simulated continuous two-dimensional habitat, which is represented by a square with a side length of *L* and containing *N* uniformly distributed individuals, the interaction radius δ is defined as 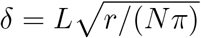.

One important consequence is that, when *D* becomes sufficiently small, the spread of the disease is no longer driven primarily by the dispersal of individuals, but instead by “hopping” between neighboring individuals with overlapping interaction circles. These “hopping” dynamics can cause the disease to fix even if individuals do not move at all, thus setting an upper bound for the fixation time in the limit *D* → 0.

We can still describe disease spread under these hopping dynamics by the diffusion model, but we need to reinterpret the diffusion term. In particular, we interpret each transmission event as a “dispersal” step of the disease from the location of the infecting individual to the location of the newly infected individual. To derive a rough estimate of the diffusion coefficient (*D*_0_) of the transmission, we consider the limit where individuals no longer move at all. Starting from the first infected individual at the center of the habitat, the disease spreads outward in a circular pattern, with the traveling wave now driven entirely by transmission events rather than by individual dispersal. The minimum speed of this traveling wave for the SI model is given by 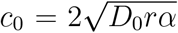 [47, 48]. One complication is that, in the hopping model, *D*_0_ varies in space because it depends on the occurrence of new transmission events. Thus, *D*_0_ approaches zero in areas where either everyone is already infected or everyone is still susceptible. It is maximal right at the wavefront, where it determines the speed of the wave.

Let us, therefore, consider an infected individual located exactly at the wavefront. If its neighbors are uniformly distributed across its interaction circle, about half of these individuals are (on average) already infected, while the others are still susceptible. The overall rate at which transmission events from the focal infected individual occur then is Pr(transmission) = *rα/*2. When a transmission event occurs, the variance of the displacement along the *x* axis is 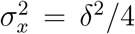. The effective diffusion coefficient of the hopping process thus results as 2*D*_0_ ≈ Pr(transmission) 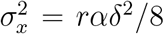. According to Equation (8), this yields a fixation time of:

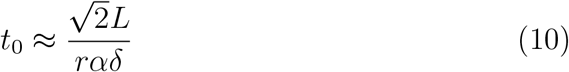

This provides an approximate upper bound for the fixation time in the limit *D* → 0, where the fixation time from dispersal alone, given by Equation (8), diverges to infinity. By comparing the fixation time estimates *t*_0_ and *t*_DIFF_, we determine *D*_*l*_, a threshold for individual dispersal below which disease spread in our individual-based simulation model is primarily driven by transmission events:

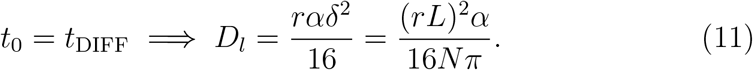

### 2.5. SI Model

To test how well our mathematical predictions describe disease spread in the individual-based simulations, we start with the SI model. Qualitatively, we expect that in the low-dispersal regime (*D < D*_*c*_ = 3.536 *×* 10^*−*6^), the disease spreads from its introduction at the center of the habitat as a circle with a steadily growing radius. By contrast, in the high-dispersal regime (*D* ≫ *D*_*c*_), the population should be sufficiently mixed so that the frequency of infected individuals increases in all areas of the habitat at a similar rate. Figure 2 confirms these qualitative predictions when comparing two simulation runs with *D* = 10^*−*6^ (left column) and *D* = 10^*−*2^ (right column).

**Figure 2:**
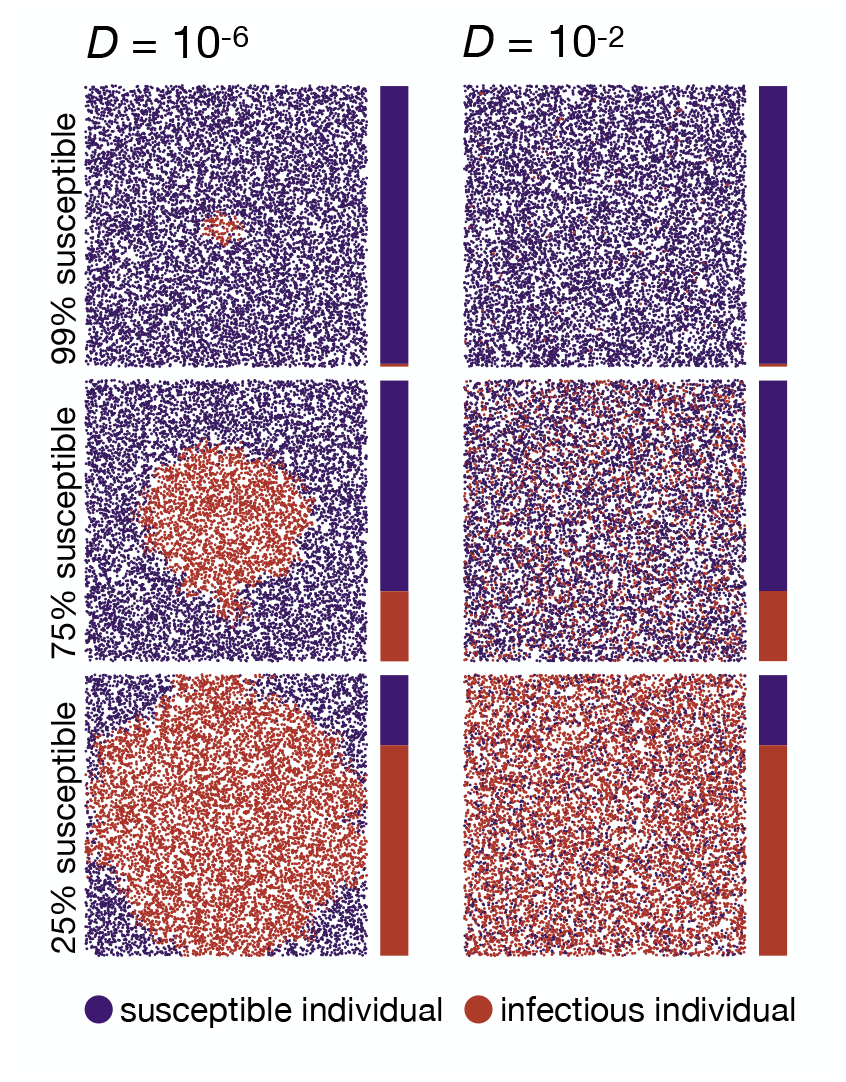
Two example runs of the SI simulation model. The left column shows a run in the low-dispersal regime (*D* = 10^*−*6^), while the right column shows a run in the high-dispersal regime (*D* = 10^*−*2^). The top, middle, and bottom plots show population snapshots taken when 99%, 75%, and 25% of the population were still susceptible. As predicted, the disease progresses in a circular pattern in the low-dispersal regime. In contrast, with high dispersal, infectious individuals are homogeneously distributed in space. Data were simulated using *N* = 10, 000 (ten times smaller than our standard model for better visualization), *α* = 0.01, *r* = 15, *L* = 1. Videos of simulation runs in the SI model can be found under https://tinyurl.com/bdddam58.

We next tested our predictions for the disease fixation time under varying dispersal rates (Figure 3A). In the high-dispersal regime, the observed fixation time *t*_sim_ is independent of *D* and well approximated by the predictions from the ODE model given in Equation (4). The time-resolved proportion of infectious individuals follows the logistic growth curve (Figure 3B, bottom panel). When *D* becomes smaller than *D*_*c*_, the fixation time starts to increase and becomes inversely proportional to *D*, as predicted by Equation (8). Once *D* becomes smaller than *D*_*l*_, the dispersal threshold below which disease spread in our individual-based simulation model is primarily driven by transmission events (*D*_*l*_ = 4.476 *×* 10^*−*8^), the fixation time approaches the prediction under the hopping dynamics derived in Equation (10), *t*_sim_ levels off, and again becomes independent of *D*.

**Figure 3:**
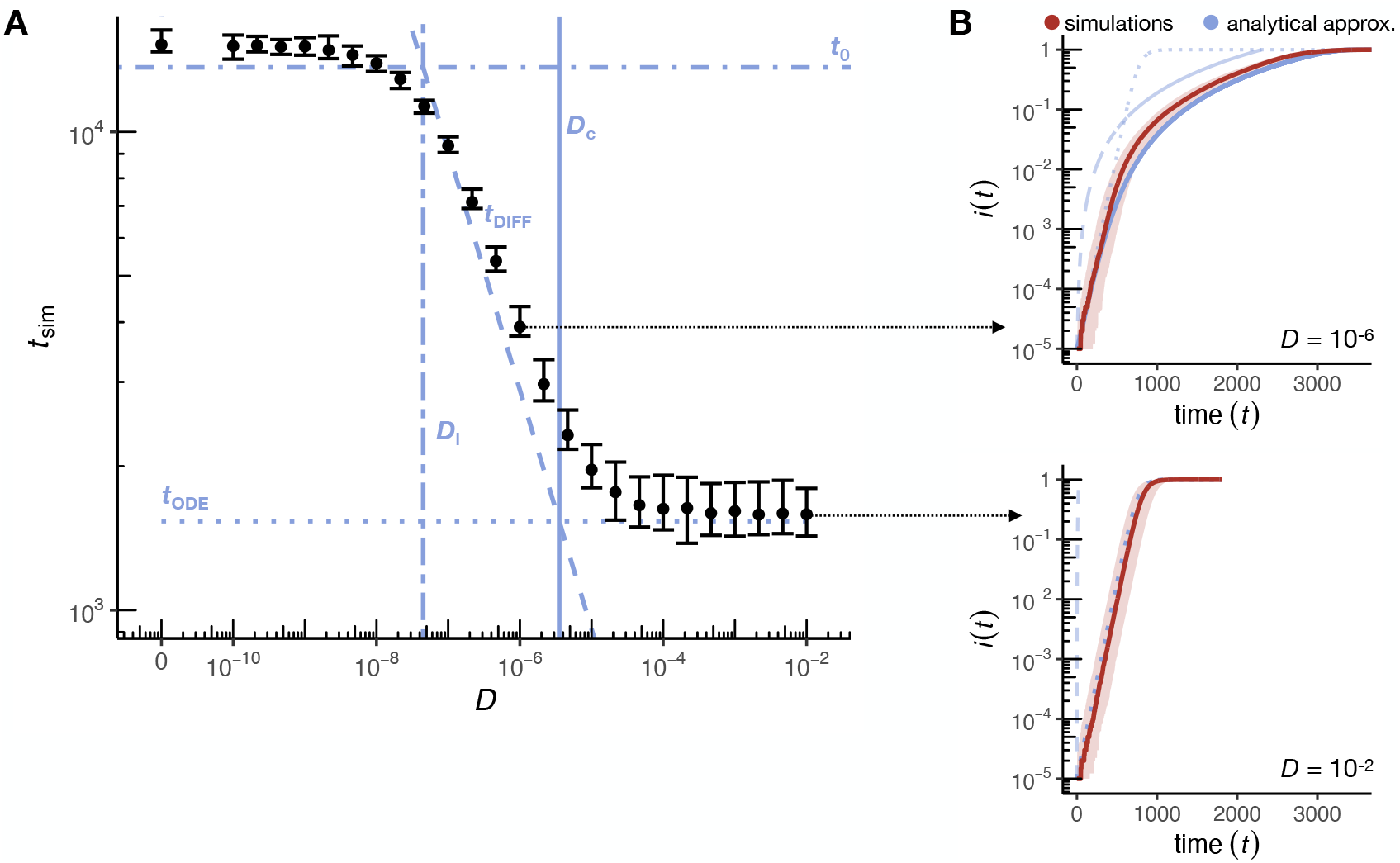
Disease dynamics in the SI model. (A) Time until disease fixation in the simulations, *t*_sim_, for varying diffusion coefficients *D*. Black dots represent median values and error bars the 2.5 and 97.5 percentiles estimated from 50 simulation runs for each *D* (for *D* = 0, the spread stagnated in 3 of the 50 runs due to small clusters of susceptible individuals with interaction areas without infectious individuals, which we excluded from the analysis). The vertical solid line indicates the critical threshold *D*_*c*_ according to Equation (9). In the high-dispersal regime (*D* ≫ *D*_*c*_), fixation times converge to the prediction of the ODE model (*t*_ODE_, dotted line), while in the limit of no dispersal (*D* → 0) they are bounded by the prediction of the hopping model (*t*_0_, dot-dashed line; *D*_*l*_ is indicated as a vertical double-dashed line). In the low-dispersal regime where *D < D*_*c*_, the disease advances as a traveling wave, and *t*_sim_ is well approximated by *t*_DIFF_ (dashed line). (B) The upper panel shows the time-resolved disease dynamics for the low-dispersal regime (*D* = 10^*−*6^). Here, the proportion of infectious individuals, *i*(*t*), increases after an initial lag phase approximately as the area of a circle whose radius expands at the predicted minimum wave speed of the diffusion model (blue dashed line). The initial lag between the expected and the observed proportion of infectious individuals is likely caused by the local establishment of the disease and an initial reduction in wave speed due to the curvature of the wavefront [8, 44]. The *i*(*t*) of the simulations is well approximated by the heuristic approximation *t*_fix_ (solid blue line) that accounts for both, the disease propagation via a constant traveling wave, and the local rise in *i*(*t*) described by the ODE model (dotted line). The bottom panel shows an example from the high-dispersal regime (*D* = 10^*−*2^). Here, the fraction of infectious individuals is well approximated by the logistic growth function of the ODE model (dotted blue line). Solid red lines represent the median across 50 simulation runs and the shaded areas represent the range between the 2.5 and 97.5 percentiles. All data were simulated using *N* = 100, 000, *α* = 0.001, *r* = 15, *L* = 1.

Overall, we observe that fixation times can differ up to one order of magnitude in our simulation model depending on the dispersal rate. This highlights the risk of underestimating the expected epidemic duration if predictions are based solely on the ODE model without accounting for population structure.

### 2.6. SIS Model

In the SI model, infected individuals do not recover from the infection, leading to an inevitable spread and eventual fixation of the disease as long as *rα >* 0. The SIS model allows infected individuals to transition from infection back to the susceptible state at rate *γ*. In a deterministic ODE model of an unstructured population, such a disease is expected to approach a constant frequency of 1−1*/R*_0_ for *R*_0_ *>* 1, where *R*_0_ = *rα/γ* represents the basic reproduction number of the disease (i.e., the average number of secondary infections caused by a single infectious individual when introduced into a completely susceptible population) [52, 4]. If *R*_0_ *<* 1, the disease is unable to become established in the population and will typically die out quickly. Much work has been done to define *R*_0_ in reaction-diffusion models, which are inherently more complex than their ODE counterparts [53]. Although the *R*_0_ definitions in reaction-diffusion models differ from the classical ODE-based definition, they can converge under certain conditions, such as when diffusion rates approach 0 [54, 53]. In this study, we use the classical ODE-based *R*_0_ as a reference point for easier comparison across different dispersal regimes.

Analogous to the SI model, one can again determine a critical threshold *D*_*c*_, below which individual dispersal is expected to affect the disease dynamics. For the SIS model, the frequency of infected individuals *i*(*t*) follows a logistic growth function with a carrying capacity of 1− 1*/R*_0_ = 1 − *γ/*(*rα*) in a homogeneously mixing population. With *I*(0) = 1,

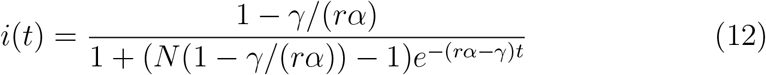

Equation 12 allows the derivation of an expected “time until constant endemic frequency” estimate for the ODE model, similar to the derivation of the “fixation time” estimate for the SI model (Equation 4). Assuming *N* ≫ 1, this “time until constant endemic frequency” can be estimated for the SIS model in a homogeneously mixing population as:

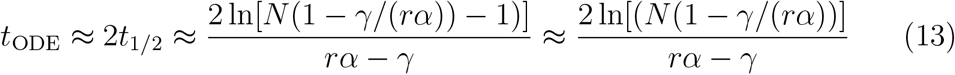

For a spatial SIS model, the recovery of infectious individuals back to the susceptible state leads to a slowing of wave propagation, with the recovery rate *γ* entering the minimum wave speed approximation as 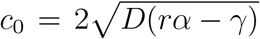 [29, 47, 48]. We thus obtain the time required for the spatial spread of the disease in the SIS model as:

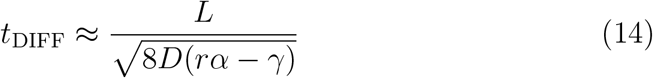

Following the lines discussed above for the SI model, we can compare the expected time to constant endemic frequency, *t*_ODE_, in the ODE model with the spread time, *t*_DIFF_, to obtain a critical value for the diffusion coefficient above which we expect a negligible effect of spatial progression on disease dynamics:

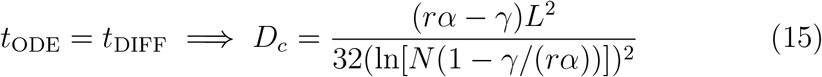

To investigate the effects of spatial structure and stochasticity on an SIS model, we modified our individual-based simulations so that infected individuals reenter the susceptible class with a probability *γ* per tick, yielding *R*_0_ = *rα/γ*. Qualitatively, we find that the disease initially still spreads in a circular fashion in the low-dispersal regime and more homogeneously in the high-dispersal regime, as in the SI model (Figure 4). However, the continuous re-entry of infectious individuals into the susceptible class breaks up this strict circular clustering more quickly in the SIS model than in the SI model. Over the course of the simulated time period, the frequency of infectious individuals converges to the expected constant endemic frequency, 1− 1*/R*_0_, in both the high-dispersal and low-dispersal regimes. We note that in our stochastic simulation model with finite population size, this constant endemic frequency is only quasi-stationary: On much longer time scales than considered here, the stochastic model will eventually reach its absorbing, disease-free state (all infectious individuals will eventually disappear from the population).

**Figure 4:**
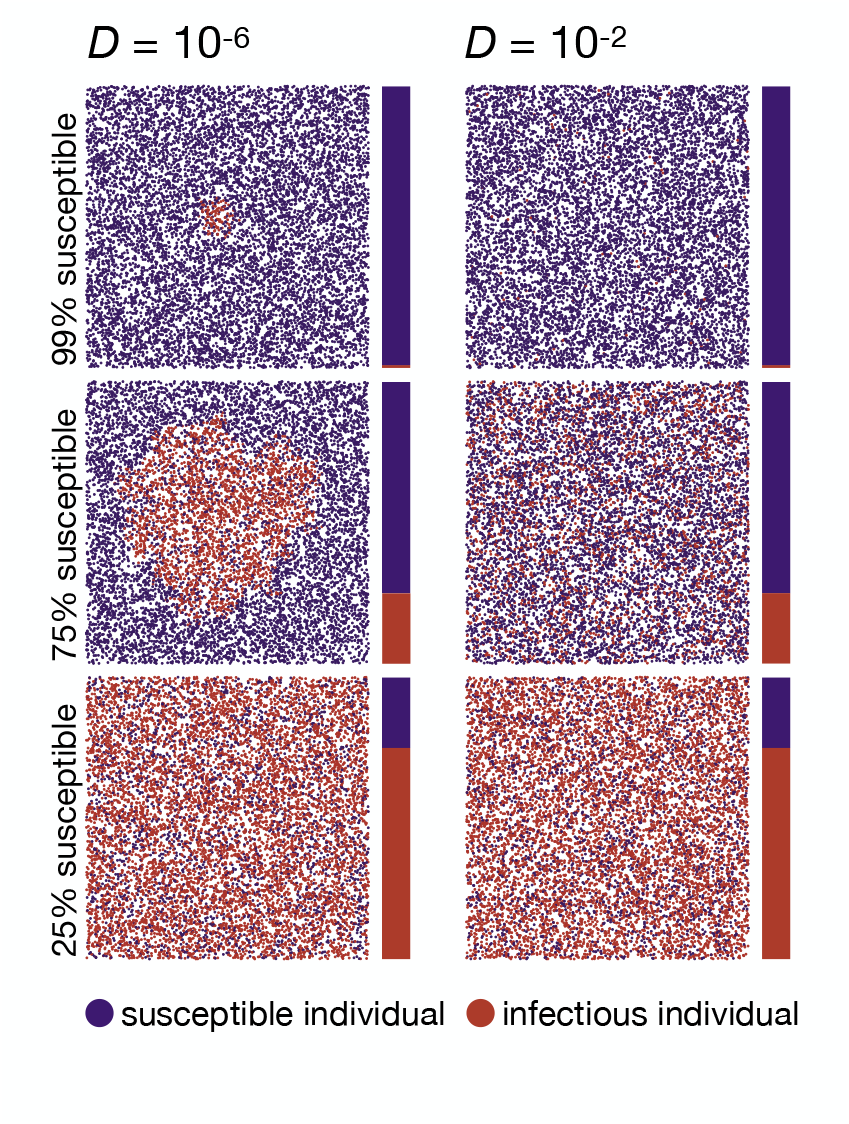
Two example runs of the SIS simulation model. The left column shows a run in the low-dispersal regime (*D* = 10^*−*6^ *< D*_*c*_), while the right column shows a run in the high-dispersal regime (*D* = 10^*−*2^). The top, middle, and bottom plots show population snapshots taken when 99%, 75%, and 25% of the population were still susceptible. As predicted, in the low-dispersal regime, the disease initially progresses in a growing circle. In contrast, in the high-dispersal regime, infectious individuals are homogeneously distributed in space (right column). In contrast to the SI model, in the SIS model, even with low dispersal, infectious individuals hardly cluster at later stages of the simulations due to the continuous recovery of infectious individuals with probability *γ* per tick. Data were simulated with *D* = [10^*−*6^; 10^*−*2^], *N* = 10, 000, *α* = 0.01, *r* = 15, *L* = 1, *γ* = 1*/*30. Videos of simulation runs in the SIS model can be found under https://tinyurl.com/bdddam58.

While a deterministic ODE model predicts that any disease with *R*_0_ *>* 1 will establish after introduction, the stochastic fluctuations in our individual-based simulation model can lead to the loss of the disease in certain simulation runs, even for relatively high *R*_0_ values (e.g., about 25% of simulation runs with *R*_0_ = 3, as shown in Figure 5A). We did not find a strong influence of the dispersal rate on the probability of disease establishment for *R*_0_ *>* 2.

**Figure 5:**
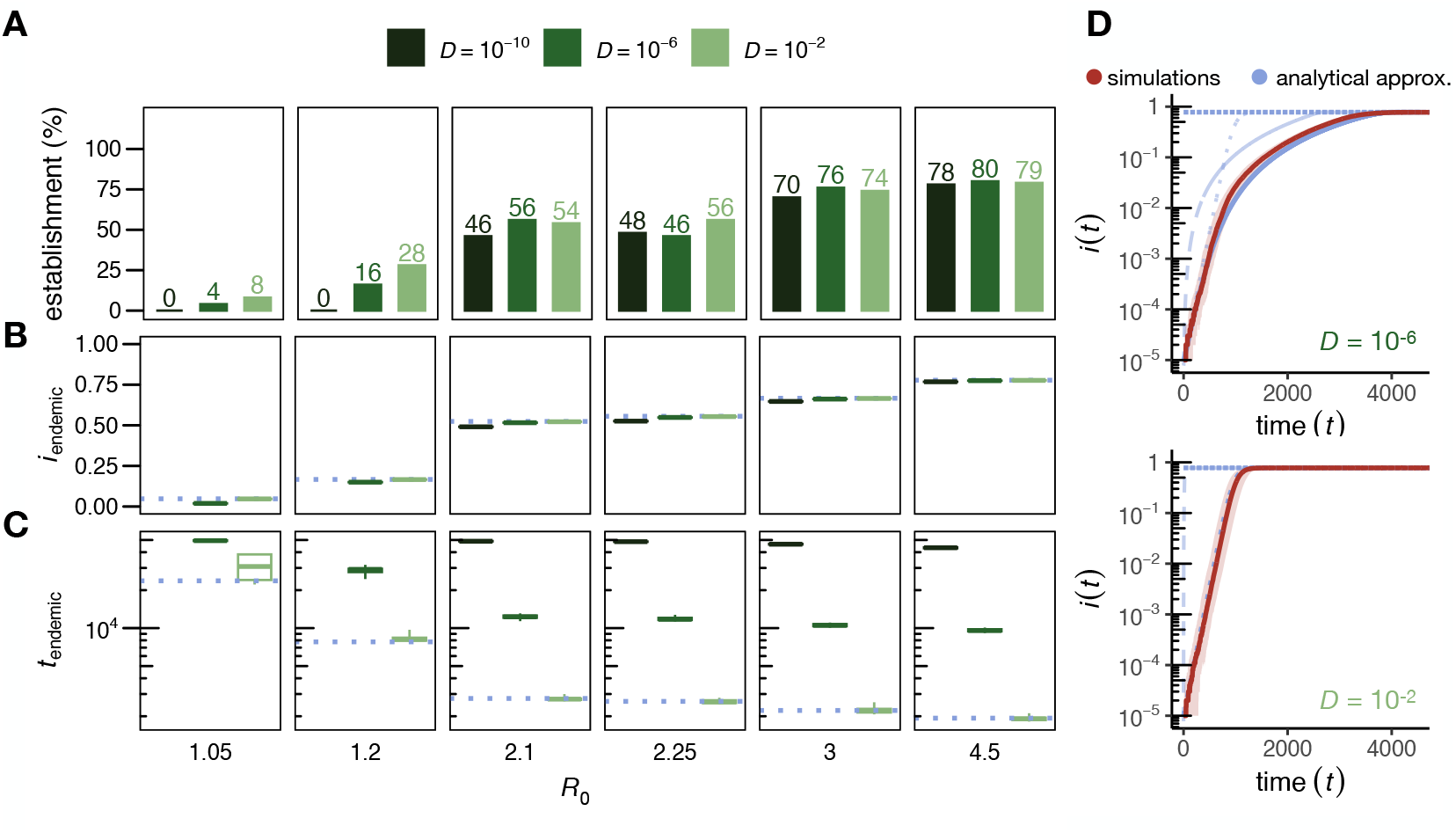
Disease dynamics in the SIS model. (A) Percentage of 50 simulation runs that reached a frequency of at least 1% infectious individuals for different *R*_0_ values (panels) and three exemplary diffusion coefficients. Only simulation runs with an *i*(*t*) of at least 1% are included in (B) to (D). (B) Observed constant endemic frequencies at the end of the simulations. Box plots are shown with whiskers indicating the most extreme values within 5 *× IQR* of the boxes, but the variance between runs was so small that they are not visible. The expected constant endemic frequency for each *R*_0_ value under an ODE model [4] is shown by a dotted blue line. Observed constant endemic frequencies were estimated by fitting simulated data to a three-parameter logistic model using the drc R package [55]. (C) Time to reach the constant endemic frequency. We defined the onset of constant *i*(*t*) as the first time the frequency of infectious individuals reaches 99.95% of the observed constant endemic frequency. The expected time of constant *i*(*t*) onset under an ODE model is depicted as a dotted blue line. (D) Time-resolved disease dynamics for *R*_0_ = 4.5 and two exemplary diffusion coefficients (upper panel: low dispersal with *D* = 10^*−*6^ *< D*_*c*_ = 2.87 *×* 10^*−*6^, lower panel: high dispersal with *D* = 10^*−*2^). In the low-dispersal regime, the proportion of infectious individuals, *i*(*t*), increases after an initial lag phase approximately as the area of a circle whose radius expands at the predicted minimum wave speed of the diffusion model (blue dashed line). The initial lag between the expected and the observed proportion of infectious individuals is likely caused by the local establishment of the disease and an initial reduction in wave speed due to the curvature of the wavefront [8, 44]. The *i*(*t*) of the simulations is well approximated by a heuristic approach *t*_fix_ (solid blue line) that accounts for both disease propagation via a constant traveling wave, and the local increase in *i*(*t*) described by the ODE model (dotted line). In the high-dispersal regime, the proportion of infectious individuals is well approximated by the expected logistic growth function of the ODE model with a carrying capacity of 1 *−* 1*/R*_0_ [4] (dotted blue line). Solid red lines represent the median proportion of infectious individuals across all simulations, and the shaded area represents the range between the 2.5 and 97.5 percentiles. All data were simulated with *N* = 100, 000, *α* = 0.001, *r* = 15, *γ* = [1*/*70, 1*/*80, 1*/*140, 1*/*150, 1*/*200, 1*/*300]. More than 99% of our simulation runs categorized as established spread through the entire habitat.

However, in the high-dispersal regime, the frequency of infected individuals approaches the expected constant endemic frequency more quickly than in the low-dispersal regime (Figure 5B-D). Overall, we observed that the rate of disease spread in our simulation model can again differ by more than an order of magnitude, depending on the dispersal rate (Figure 5C).

### 2.7. SIR Model

In the SIS model, individuals transition between the susceptible and infectious states but do not become resistant to the disease. In contrast, in the SIR model, infected individuals acquire permanent resistance and transition to a “recovered state” that cannot be left under the dynamic. In the unstructured SIR model, a disease can invade a fully susceptible population whenever its basic reproduction number *R*_0_ = (*rα*)*/γ* is greater than 1 [4]. In such cases, *i*(*t*) first increases to a maximum value *i*_max_, then decreases over time, approaching zero as *t* → ∞. In contrast to the SI and SIS models, there is no exact analytical solution for *i*(*t*) and approximations are often complex [56, 57]. However, the maximum incidence *i*_max_ can be derived. With *I*(0) = 1 and *S*(0) = *N −* 1, [5]:

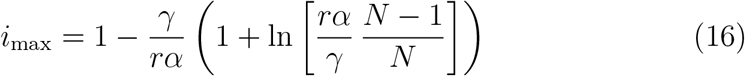

For the time *t*_ODE_ to reach this maximum, *i*(*t*_ODE_) = *i*_max_, a useful approximation has been derived (Eq. 7a of Turkyilmazoglu (2021) [58]):

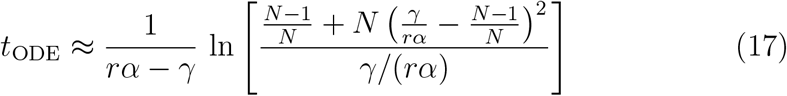

Finally, the proportion of all individuals infected at any time during the epidemic, the so-called “final epidemic size”, *e*_final_, can be obtained by solving the transcendental equation [3],

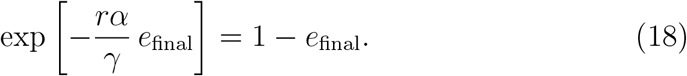

As in the SI and SIS models, we can derive a critical threshold *D*_*c*_ for the SIR model, below which individual dispersal affects disease dynamics. We use that the speed of the infection wave in the spatial SIR model is the same as in the spatial SIS model, 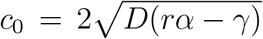 [29, 47, 48]. Thus, the expected time *t*_DIFF_ for wave propagation in the spatial SIR model is also equal to the time in the spatial SIS model (equation 14). Since the infection wave is closely followed by a wave of recovered individuals who cannot be reinfected, the proportion of infectious individuals, *i*(*t*), expands approximately as the area of a circular ring of increasing radius and reaches its maximum when this ring reaches the corners of the square habitat patch. As for the SI and SIS models, we obtain a critical value *D*_*c*_ of the diffusion coefficient by comparing the expected times required for local growth, *t*_ODE_ and for spatial spread, *t*_DIFF_, respectively, to reach the maximum *i*_max_ of *i*(*t*). Specifically, we expect individual dispersal to influence disease dynamics in the spatial SIR model for *D < D*_*c*_ with

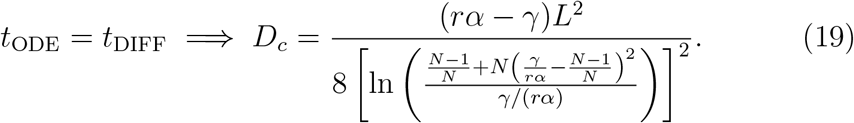

To study the SIR model, we modified our simulations of the SIS model so that infected individuals transition to the recovered class with a constant probability *γ* per tick, rather than returning to the susceptible class. This modification does not change the basic reproduction number of the disease, which remains at *R*_0_ = *rα/γ*. Qualitatively, we observed that, similar to the SI and SIS models, the high-dispersal regime closely approximates a homogeneously mixed population (Figure 6), whereas in the low-dispersal regime, the disease spreads in a circular pattern across space, as theoretically predicted [59]. However, the traveling wave by which infections spread is now closely followed by another wave describing the emergence of recovered individuals.

**Figure 6:**
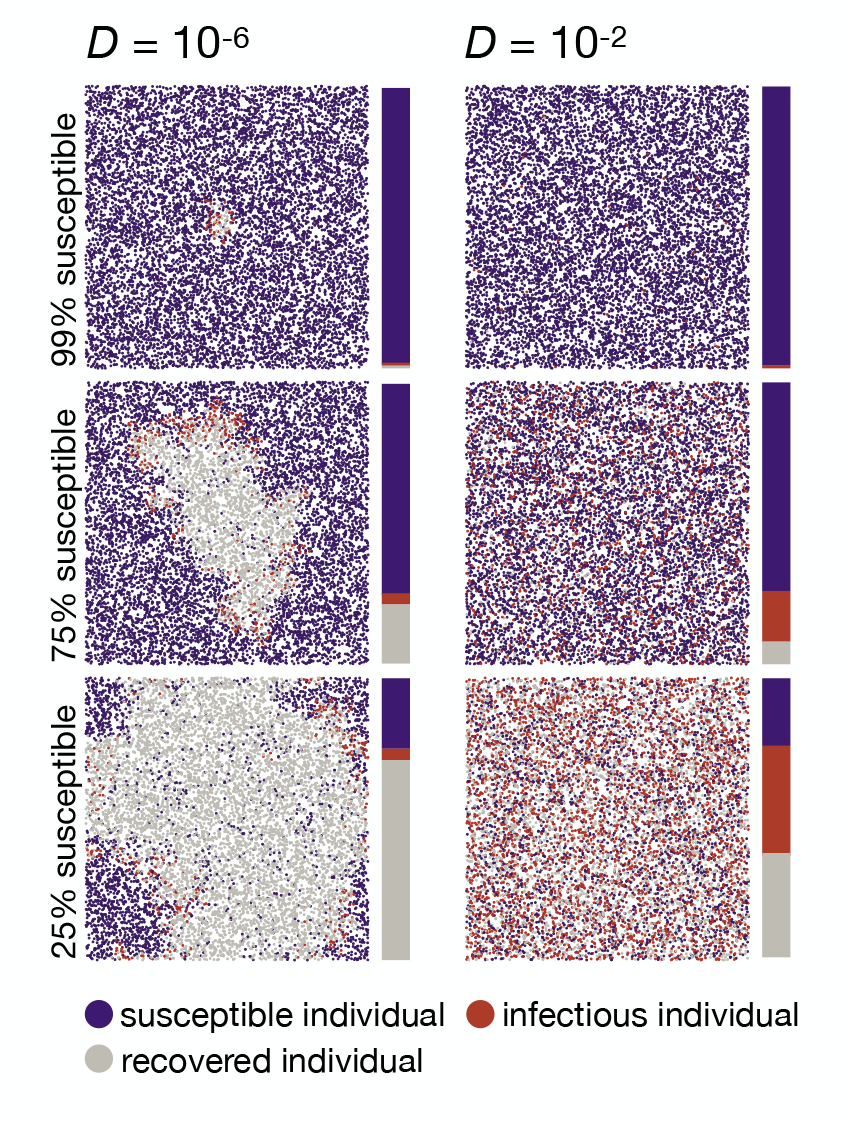
Two example runs of the SIR simulation model. The left and right columns show runs in the low-dispersal (*D* = 10^*−*6^ *< D*_*c*_) and high-dispersal (*D* = 10^*−*2^) regimes, respectively. The top, middle, and bottom plots show population snapshots taken when 99%, 75%, and 25% of the population were still susceptible. As predicted, in the low-dispersal regime, the disease progresses approximately in a growing circle, with infectious individuals clustered at the wave front, and recovered individuals clustered around the disease origin in the center of the area. In contrast, in the high-dispersal regime susceptible, infectious, and recovered individuals are homogeneously distributed in space (right column). Data were simulated with *D* = [10^*−*6^; 10^*−*2^], *N* = 10, 000, *α* = 0.01, *r* = 15, *L* = 1, *γ* = 1*/*25. Videos of simulation runs in the SIR model can be found under https://tinyurl.com/bdddam58.

The clustering of recovered individuals close to the wavefront of infectious individuals has several implications for disease dynamics: First, similar to the SIS model, for *R*_0_ *<* 2 there is a higher probability that the disease will be eradicated before it can establish in the low-dispersal regime. This results in a lower probability of establishment for smaller diffusion coefficients. For very weak diffusion (*D* = 10^*−*10^), the establishment probability remains low even until *R*_0_ exceeds 3 (Figure 7A). Second, as previously demonstrated [10, 9], higher values of *R*_0_ are required in the low-dispersal regime to approach the theoretically expected final epidemic size (Figure 7B). If dispersal is limited, the clustering of recovered individuals near the wavefront can lead to isolated ‘pockets’ of susceptible individuals without contact with still infectious individuals. This phenomenon can contribute significantly to reducing the final epidemic size, as illustrated in Figure 6 for *D* = 10^*−*6^, where distinct clusters of susceptible individuals surrounded by recovered individuals are observed. Third, because new infections occur only at the wavefront in the low-dispersal regime, the maximum incidence (*i*_max_) is smaller than *i*_max_ in the high-dispersal regime, which agrees well with the analytical expectations from the ODE model (Figure 7C). Analogous to the SI and SIS models, reduced dispersal slows down disease spread. Simulations with a diffusion coefficient of *D* = 10^*−*6^ take almost an order of magnitude longer to reach their respective *i*_max_ compared to simulations with *D* = 10^*−*2^ (Figure 7D). Finally, the time-resolved frequency of infectious individuals, *i*(*t*), resembles a bell-shaped curve in the high-dispersal regime, as predicted by the ODE model [4] and the observed peak times agree well with the peak time approximations for the ODE model [58]. In the low-dispersal regime, on the other hand, *i*(*t*) shows nearly linear growth until late in the simulation, as predicted by a circular wave progressing at constant velocity [29, 44, 47, 48] (Figure 7D-E).

**Figure 7:**
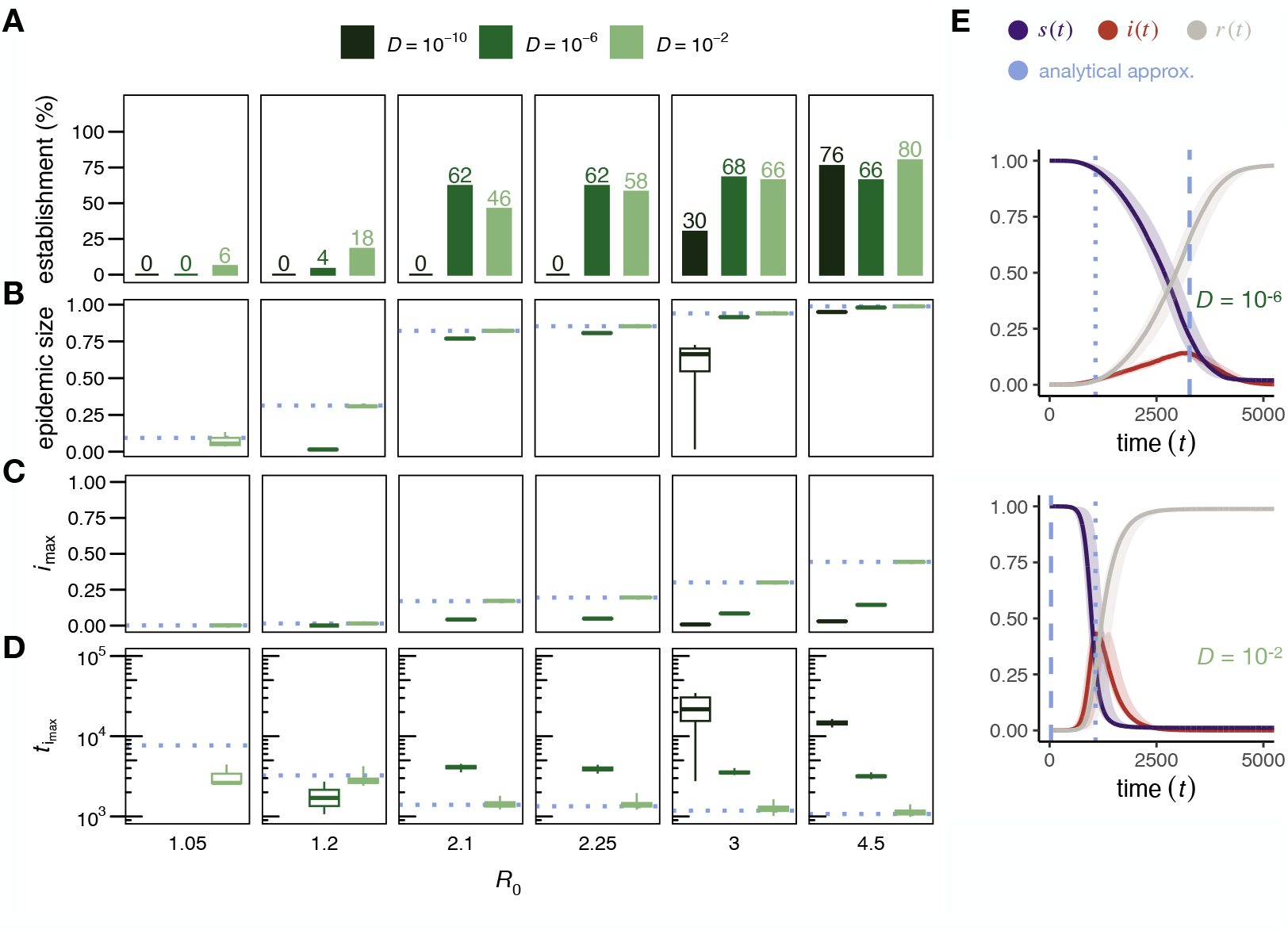
Disease dynamics in the SIR model. (A) Percentage of 50 simulation runs in which the disease reached an epidemic size of at least 1% for different values of *R*_0_ (panels) and three exemplary diffusion coefficients (for *D* = 10^*−*10^ and *R*_0_ = 3, the spread stagnated in 3 of the 50 runs, which were excluded from the analysis). Disease establishment is less likely in the low-dispersal regime for *R*_0_ *<* 2. Only simulation runs with an epidemic size of at least 1% are included in (B) to (E). (B) Epidemic size (box plots are shown with whiskers indicating the most extreme values within 5 *× IQR* of the boxes). Simulations in the low-dispersal regime require higher *R*_0_ values to approach the theoretically predicted epidemic size under the ODE model. (C) Maximum incidence *i*_max_ (the variance between runs was so small that the whiskers are not visible). The expected *i*_max_ [5] for each *R*_0_ value under an ODE model is shown by the dotted blue line. Lower dispersal results in smaller *i*_max_ values. (D) Peak Times. Simulations in the low-dispersal regime can take an order of magnitude longer to reach *i*_max_ than in the high-dispersal regime. The peak time approximation under an ODE model [58] is shown as dotted blue line. (E) Time-resolved disease dynamics for *R*_0_ = 4.5 and two exemplary diffusion coefficients (upper panel: low-dispersal with *D* = 10^*−*6^ *< D*_*c*_ = 9.31 × 10^*−*6^, lower panel: high-dispersal with *D* = 10^*−*2^). In the low-dispersal regime, the spatial progression of *i*(*t*) can be described as a circular ring whose radius expands with the predicted minimum wave speed and reaches its maximum around the expected peak time in a spatial SIR model (dashed blue line). In the high-dispersal regime, on the other hand, *i*(*t*) increases rapidly and reaches its maximum around the expected peak time in an ODE model [58] (dotted blue line). The solid lines represent the median proportions of susceptible (blue), infectious (red), and recovered (grey) individuals across all 50 simulations, and the shaded areas represent the ranges between the 2.5 and 97.5 percentiles. All data were simulated with *N* = 100, 000, *α* = 0.001, *r* = 15, *γ* = [1*/*70, 1*/*80, 1*/*140, 1*/*150, 1*/*200, 1*/*300]. More than 99% of our simulation runs categorized as established spread through the entire habitat.

## 3. Discussion

In this study, we investigated the conditions under which it is essential to incorporate spatial structure into epidemiological models in order to obtain accurate predictions of disease spread in time and space. We established a critical threshold *D*_*c*_ for the diffusion coefficient, below which disease transmission dynamics are expected to exhibit substantial spatial heterogeneity, while for *D* ≫ *D*_*c*_ the assumption of homogeneous mixing remains adequate. To validate our analytical results, we performed individual-based simulations on a continuous two-dimensional landscape. Furthermore, we investigated the impact of continuous spatial structure on key epidemiological parameters, including disease establishment probability, maximum incidence, peak time, and final epidemic size.

The use of reaction-diffusion models in epidemiology is attractive because of their ability to approximate epidemic states as individuals move and interact in a spatial domain. They often provide useful estimates for the expected speed of disease propagation based on measurable life-history traits, such as individual dispersal, average contact rate, and infection risk [36]. In this study, we focus on the classical Fisher-KPP equation to model diffusive disease transmission because of its simplicity and accessibility to applied modelers. Reaction-diffusion systems with alternative reaction terms and different propagation dynamics have been studied [60] and exact closed-form solutions for traveling wave speeds are available in the literature (e.g., Chen *et al*. 2012 [61]). Exploring alternative reaction terms could lead to broader insights into epidemic spread, but such directions are beyond the scope of this paper.

The Fisher-KPP model assumes that disease transmission is primarily local. Long-range dispersal is rare or absent [25, 62]. In addition, distributed-infectives models, as studied here, assume that dispersal is uniform throughout the population, ignoring variations in movement patterns among individuals [32]. In reality, individual movement patterns can vary considerably, both in frequency and in direction or distance [63, 64]. All of these processes can significantly affect the disease dynamics and limit the accuracy of a traveling wave model. For example, an asymmetric dispersal kernel, unlike the radially symmetric one used here, produces an irregular wavefront [8]. Similarly, frequent long-range dispersal, as observed in cases such as wind-dispersed plant pathogens or avian influenza [44, 27, 33, 65, 25], can cause the disease to spread from multiple locations. In such cases, reaction-diffusion models that assume an exponentially bounded dispersal kernel [66, 65, 38] underestimate the rate of spread [44, 65, 67, 33], complicating efforts to predict and manage outbreaks [68].

Furthermore, our model assumes constant rates of disease transmission and recovery. In real populations, however, these parameters are often variable. For example, the contact rate *r* may differ for individuals of different ages [36, 69, 70] and is typically influenced by landscape features such as major roads, mountains, or rivers [36, 34]. It also varies with population density or changes in behavior [71]. The probability of disease establishment *α* is often strongly influenced by environmental factors such as temperature, relative humidity, and the level of urbanization [72]. All of these factors can play a critical role in shaping the disease dynamics [73], yet they are not explicitly included in our current modeling framework.

Finally, in the Fisher-KPP framework, space is a continuous, unbounded domain. This ignores edge effects that exist in real habitats with boundaries [28, 29, 5, 51]. To focus on the effect of individual dispersal on disease spread, without confounding by edge effects, we implemented periodic boundary conditions in our individual-based simulation framework. Reflecting boundaries may better simulate real-world habitats where physical barriers constrain disease spread. However, any resulting boundary effects are highly situational (e.g., depending on the shape of the habitat and the location of the disease outbreak), but generally small in a sufficiently large habitat.

Despite these limitations, there are biological systems that satisfy the assumptions of a reaction-diffusion model with a constant traveling wave. For example, Noble’s analysis of the spread of the Black Death across 14th-century Europe used a reaction-diffusion model, with the predicted spread patterns aligning well with the historical records [32]. With parameters *rα* ≈ 1-4 per year and *D* ≈ 25, 900 km^2^ per year, the distributed-infective process produced a wave “width” of 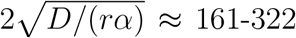 km: small relative to the studied region (Messina to Oslo is approximately 3,200 km). Furthermore, long-range dispersal was likely very rare in medieval Europe, supporting the suitability of a constant traveling wave model for studying this particular epidemic. Murray & Brown applied a reaction-diffusion model to potential rabies transmission among foxes in the UK [74]. Rabies is transmitted by close contact, and foxes are highly territorial, which limits individual dispersal. The model’s *rα* and *D* values indicated a small wave width relative to the habitat studied (southern England), with predictions matching rabies spread patterns observed in mainland Europe [74].

While it was reasonable to ignore long-range dispersal in the aforementioned studies, the increasingly interconnected nature of modern human populations requires a model that captures both short- and long-range dispersal [75, 65, 38]. Integrating these dispersal scales is essential for accurate modeling of human disease spread [12]. For recent human epidemics, Brockmann *et al*. introduced an “effective distance” measure that accounts for varying levels of connectivity between communities, ultimately allowing the application of relatively simple diffusion models to study complex disease dynamics [12]. However, such connectivity data are rarely available for non-human species, and factors such as behavior, social structures, and environmental conditions can further complicate movement patterns [38, 23, 36, 12, 76]. For complex movement patterns, network models or individual-based simulations may be more appropriate, although these increasingly complex models also limit the ability to draw general conclusions [38, 36].

The critical threshold *D*_*c*_ (Equation 9) provides a simple back-of-the-envelope calculation of the amount of individual dispersal required to noticeably slow the disease spread. For example, consider a theoretical measles outbreak in a fully susceptible population. Measles, with up to 90% transmission among close contacts (*α* = 0.9) [77], and assuming four close contacts per day (*r* = 4) [78], in a population with 979 individuals per km^2^ (i.e., the average population density in urbanized areas of the U.S.) [79], results in *D*_*c*_ = 0.002 km^2^ per day. In an area of *L* = 1 km, this means that for limited dispersal to significantly slow down the spread of measles, the mean standard deviation of the spatial displacement of individuals in each of the *x* and *y* coordinates must be less than 69 meters per day. This toy example shows that spatial dynamics can be neglected for highly contagious diseases over small areas [50], but must be considered for larger communities, such as cities [80, 37].

Our individual-based simulations highlighted important differences and complexities that arise when simulating a continuous reaction-diffusion process. In our discrete framework, we defined contacts between individuals based on an interaction radius parameter, allowing disease to spread by “hopping” between neighboring individuals, even when individuals are stationary. This mechanism imposes an upper limit on the fixation time in our individual-based simulations, which may not be immediately apparent in other disease models. While a *k*-nearest-neighbor approach could mitigate the influence of interaction radius, caution is warranted when using small *k* values (e.g., *k* = 1) in combination with low dispersal. Such configurations can substantially slow disease spread and, in some cases, result in coexistence between susceptible and infectious individuals: a phenomenon not predicted by continuous SI or SIR models. Durrett and Levin observed conceptually similar outcomes in their seminal work studying species interactions within spatially distributed populations [81]. They noted that while continuous models predict the extinction of both species when the two species compete but one cannot survive without the other (analogous to our case where infectious individuals require susceptibles to propagate the infection), discrete models allow for coexistence, particularly in simulation scenarios with low population densities (and thus, reduced inter-species contact rates) [81].

In conclusion, our study highlights the need for careful interpretation and understanding of spatial factors in epidemiological dynamics, while also demonstrating the complexities and design choices that arise in modeling such dynamics.

## 4. Data Accessibility

The individual-based SI, SIS, and SIR disease transmission models are implemented in the open-source software SLiM (version 4.0) [40] and are available on GitHub under https://github.com/AnnaMariaL/SpatialDiseaseSim. Videos demonstrating disease spread in structured and unstructured populations are available on YouTube under https://tinyurl.com/bdddam58.

## 5. Acknowledgements

We thank Brandon D. Hollingsworth for valuable feedback and discussion, and Benjamin C. Haller for helpful advice on the implementation of the individual-based simulation framework. This project has received funding from the European Union’s Horizon2020 research and innovation program under the Marie Sk lodowska-Curie grant agreement No. 101025586. PWM was supported by the National Institutes of Health under Award Number R35GM152242.

## References

[1] W. O. Kermack, A. G. McKendrick, A contribution to the mathematical theory of epidemics, Proceedings of the Royal Society of London. Series A, Containing Papers of a Mathematical and Physical Character 115 (772) (1927) 700–721. doi:10.1098/rspa.1927.0118.

[2] R. M. Anderson, R. M. May, Infectious diseases of humans: dynamics and control, Oxford University Press, 1991. doi:10.1093/oso/9780198545996.001.0001.

[3] J. D. Murray, Mathematical Biology I. An Introduction, 3rd Edition, Vol. 17 of Interdisciplinary Applied Mathematics, Springer, 2002. doi: 10.1007/b98868.

[4] H. W. Hethcote, Three Basic Epidemiological Models, Springer Berlin Heidelberg, Berlin, Heidelberg, 1989, Ch. 1, pp. 119–144. doi:10.1007/978-3-642-61317-3_5.

[5] H. W. Hethcote, The mathematics of infectious diseases, SIAM Review 42 (4) (2000) 599–653. doi:10.1137/S0036144500371907.

[6] M. M. López-Flores, D. Marchesin, V. Matos, S. Schecter, Differential Equation Models in Epidemiology, IMPA, 2021. URL https://schecter.math.ncsu.edu/33CBM09-eBook.pdf

[7] N. C. Grassly, C. Fraser, Mathematical models of infectious disease transmission, Nature Reviews Microbiology 6 (6) (2008) 477–487. doi: 10.1038/nrmicro1845.

[8] M. Lewis, P. Kareiva, Allee dynamics and the spread of invading organisms, Theoretical Population Biology 43 (2) (1993) 141–158. doi: 10.1006/tpbi.1993.1007.

[9] D. Mollison, Dependence of epidemic and population velocities on basic parameters, Mathematical Biosciences 107 (2) (1991) 255–287. doi: 10.1016/0025-5564(91)90009-8.

[10] B. T. Grenfell, A. P. Dobson (Eds.), The Ecology of Infectious Diseases in Natural Populations, Cambridge University Press, Cambridge, 1995. doi:10.1017/CBO9780511629396.

[11] S. Riley, Large-scale spatial-transmission models of infectious disease, Science 316 (5829) (2007) 1298–1301. doi:10.1126/science.1134695.

[12] D. Brockmann, D. Helbing, The hidden geometry of complex, network-driven contagion phenomena, Science 342 (6164) (2013) 1337–1342. doi: 10.1126/science.1245200.

[13] C. Rozins, M. J. Silk, D. P. Croft, R. J. Delahay, D. J. Hodgson, R. A. McDonald, N. Weber, M. Boots, Social structure contains epidemics and regulates individual roles in disease transmission in a group-living mammal, Ecology and Evolution 8 (23) (2018) 12044–12055. doi:10.1002/ece3.4664.

[14] C. Buckee, A. Noor, L. Sattenspiel, Thinking clearly about social aspects of infectious disease transmission, Nature 595 (7866) (2021) 205–213. doi:10.1038/s41586-021-03694-x.

[15] V. Capasso, Mathematical structures of Epidemic Systems, Springer Berlin Heidelberg, 1993. doi:10.1007/978-3-540-70514-7.

[16] S. Riley, K. Eames, V. Isham, D. Mollison, P. Trapman, Five challenges for spatial epidemic models, Epidemics 10 (2015) 68–71, challenges in Modelling Infectious Disease Dynamics. doi:10.1016/j.epidem.2014.07.001.

[17] F. Brauer, C. Castillo-Chavez, Z. Feng, Spatial Structure in Disease Transmission Models, Springer New York, New York, NY, 2019, Ch. 14, pp. 457–476. doi:10.1007/978-1-4939-9828-9_14.

[18] A. Lajmanovich, J. A. Yorke, A deterministic model for gonorrhea in a nonhomogeneous population, Mathematical Biosciences 28 (3) (1976) 221–236. doi:10.1016/0025-5564(76)90125-5.

[19] V. Belik, T. Geisel, D. Brockmann, Natural human mobility patterns and spatial spread of infectious diseases, Phys. Rev. X 1 (2011) 011001. doi:10.1103/PhysRevX.1.011001.

[20] A. Wesolowski, T. Qureshi, M. F. Boni, P. R. Sundsøy, M. A. Johansson, S. B. Rasheed, K. Engø-Monsen, C. O. Buckee, Impact of human mobility on the emergence of dengue epidemics in pakistan, Proceedings of the National Academy of Sciences 112 (38) (2015) 11887–11892. doi:10.1073/pnas.1504964112.

[21] J. I. Blanford, Z. Huang, A. Savelyev, A. M. MacEachren, Geo-located tweets. enhancing mobility maps and capturing cross-border movement, PLOS ONE 10 (6) (2015) 1–16. doi:10.1371/journal.pone.0129202.

[22] T. A. Perkins, R. C. Reiner, Jr., G. España, Q. A. ten Bosch, A. Verma, K. A. Liebman, V. A. Paz-Soldan, J. P. Elder, A. C. Morrison, S. T. Stoddard, U. Kitron, G. M. Vazquez-Prokopec, T. W. Scott, D. L. Smith, An agent-based model of dengue virus transmission shows how uncertainty about breakthrough infections influences vaccination impact projections, PLOS Computational Biology 15 (3) (2019) 1–32. doi:10.1371/journal.pcbi.1006710.

[23] R. C. Reiner, S. T. Stoddard, T. W. Scott, Socially structured human movement shapes dengue transmission despite the diffusive effect of mosquito dispersal, Epidemics 6 (2014) 30–36. doi:10.1016/j.epidem.2013.12.003.

[24] N. M. Ferguson, C. A. Donnelly, R. M. Anderson, Transmission intensity and impact of control policies on the foot and mouth epidemic in great britain, Nature 413 (6855) (2001) 542–548. doi:10.1038/35097116.

[25] M. J. Keeling, K. T. D. Eames, Networks and epidemic models, Journal of The Royal Society Interface 2 (4) (2005) 295–307. doi:10.1098/rsif.2005.0051.

[26] J. Rushmore, D. Caillaud, L. Matamba, R. M. Stumpf, S. P. Borgatti, S. Altizer, Social network analysis of wild chimpanzees provides insights for predicting infectious disease risk, Journal of Animal Ecology 82 (5) (2013) 976–986. doi:10.1111/1365-2656.12088.

[27] N. Shigesada, K. Kawasaki, Biological Invasions: Theory and Practice, Oxford University Press, 1997. doi:10.1093/oso/9780198548522.001.0001.

[28] H. Hotelling, A mathematical theory of migration, University of Washington, republished in Environment and Planning A 10 1223–1239 (1921). doi:10.1068/a101223.

[29] R. A. Fisher, The wave of advance of advantageous genes, Annals of Eugenics 7 (4) (1937) 355–369. doi:10.1111/j.1469-1809.1937.tb02153.x.

[30] A. Kolmogorov, I. Petrovskii, N. Piscunov, A study of the equation of diffusion with increase in the quantity of matter, and its application to a biological problem, Byul. Moskovskogo Gos. Univ. 1 (6) (1937) 1–25.

[31] T. Wang, Dynamics of an epidemic model with spatial diffusion, Physica A 409 (2014) 119–129. doi:10.1016/j.physa.2014.04.028.

[32] J. V. Noble, Geographic and temporal development of plagues, Nature 250 (5469) (1974) 726–729. doi:10.1038/250726a0.

[33] C. C. Mundt, K. E. Sackett, L. D. Wallace, C. Cowger, J. P. Dudley, Long-distance dispersal and accelerating waves of disease: Empirical relationships., The American Naturalist 173 (4) (2009) 456–466. doi: 10.1086/597220.

[34] D. L. Smith, B. Lucey, L. A. Waller, J. E. Childs, L. A. Real, Predicting the spatial dynamics of rabies epidemics on heterogeneous landscapes, Proceedings of the National Academy of Sciences 99 (6) (2002) 3668–3672. doi:10.1073/pnas.042400799.

[35] J. Novembre, A. P. Galvani, M. Slatkin, The geographic spread of the ccr5 Δ32 hiv-resistance allele, PLOS Biology 3 (11) (10 2005). doi: 10.1371/journal.pbio.0030339.

[36] A. Hastings, K. Cuddington, K. F. Davies, C. J. Dugaw, S. Elmendorf Freestone, S. Harrison, M. Holland, J. Lambrinos, U. Malvadkar, A. Melbourne, K. Moore, C. Taylor, D. Thomson, The spatial spread of invasions: new developments in theory and evidence, Ecology Letters 8 (1) (2005) 91–101. doi:10.1111/j.1461-0248.2004.00687.x.

[37] D. Chen, Modeling the Spread of Infectious Diseases: A Review, John Wiley and Sons, Ltd, 2014, Ch. 2, pp. 19–42. doi:10.1002/9781118630013.ch2.

[38] M. C. Steiner, J. Novembre, Population genetic models for the spatial spread of adaptive variants: A review in light of SARS-CoV-2 evolution, PLOS Genetics 18 (9) (2022) 1–22. doi:10.1371/journal.pgen.1010391.

[39] D. Mollison, The rate of spatial propagation of simple epidemics, in: Proceedings of the Sixth Berkeley Symposium on Mathematical Statistics and Probability (Univ.California, Berkeley, Calif., 1970/1971), Vol. III: Probability theory, Univ. California Press, Berkeley, Calif., 1972, pp. 579–614. URL http://projecteuclid.org/euclid.bsmsp/1200514358

[40] B. C. Haller, P. W. Messer, Slim 4: Multispecies eco-evolutionary modeling, The American Naturalist 201 (5) (2023) E127–E139. doi: 10.1086/723601.

[41] A. Tsoularis, J. Wallace, Analysis of logistic growth models, Mathematical Biosciences 179 (1) (2002) 21–55. doi:10.1016/S0025-5564(02)00096-2.

[42] H. Scherm, On the velocity of epidemic waves in model plant disease epidemics, Ecological Modelling 87 (1) (1996) 217–222. doi:10.1016/0304-3800(95)00030-5.

[43] A. Hastings, Models of spatial spread: A synthesis, Biological Conservation 78 (1) (1996) 143–148. doi:10.1016/0006-3207(96)00023-7.

[44] N. Shigesada, K. Kawasaki, Y. Takeda, Modeling stratified diffusion in biological invasions, The American Naturalist 146 (08 1995). doi: 10.1086/285796.

[45] N. Tkachenko, J. D. Weissmann, W. P. Petersen, G. Lake, C. P. E. Zollikofer, S. Callegari, Individual-based modelling of population growth and diffusion in discrete time, PLOS ONE 12 (4) (2017) 1–22. doi: 10.1371/journal.pone.0176101.

[46] P. C. Fife, Mathematical aspects of reacting and diffusing systems, Springer Berlin, Heidelberg, 1979. doi:10.1007/978-3-642-93111-6.

[47] A. Källén, Thresholds and travelling waves in an epidemic model for rabies, Nonlinear Analysis: Theory, Methods and Applications 8 (8) (1984) 851–856. doi:10.1016/0362-546X(84)90107-X.

[48] H. Wang, X.-S. Wang, Traveling Wave Phenomena in a Kermack– McKendrick SIR Model, Journal of Dynamics and Differential Equations 28 (1) (2016) 143–166. doi:10.1007/s10884-015-9506-2.

[49] Q. Zhuang, J. Wang, A spatial epidemic model with a moving boundary, Infectious Disease Modelling 6 (2021) 1046–1060. doi:10.1016/j.idm.2021.08.005.

[50] M. Kot, Elements of Mathematical Ecology, Cambridge University Press, West Nyack, 2001. doi:10.1017/CBO9780511608520.

[51] R. Mazzucco, M. Doebeli, U. Dieckmann, The influence of habitat boundaries on evolutionary branching along environmental gradients, Evolutionary Ecology 32 (12 2018). doi:10.1007/s10682-018-9956-1.

[52] K. Dietz, The estimation of the basic reproduction number for infectious diseases, Statistical Methods in Medical Research 2 (1) (1993) 23–41. doi:10.1177/096228029300200103.

[53] C. Yang, J. Wang, Computation of the basic reproduction numbers for reaction-diffusion epidemic models, Mathematical Biosciences and Engineering 20 (8) (2023) 15201–15218. doi:10.3934/mbe.2023680.

[54] S. Chen, J. Shi, Asymptotic profiles of basic reproduction number for epidemic spreading in heterogeneous environment, SIAM Journal on Applied Mathematics 80 (3) (2020) 1247–1271. doi:10.1137/19M1289078.

[55] C. Ritz, F. Baty, J. C. Streibig, D. Gerhard, Dose-response analysis using r, PLOS ONE 10 (12) (2016) 1–13. doi:10.1371/journal.pone.0146021.

[56] L. Bougoffa, S. Bougouffa, A. Khanfer, Approximate and Parametric Solutions to SIR Epidemic Model, Axioms 13 (3) (2024). doi:10.3390/axioms13030201.

[57] N. S. Barlow, S. J. Weinstein, Accurate closed-form solution of the SIR epidemic model, Physica D: Nonlinear Phenomena 408 (2020) 132540. doi:10.1016/j.physd.2020.132540.

[58] M. Turkyilmazoglu, Explicit formulae for the peak time of an epidemic from the SIR model, Physica D: Nonlinear Phenomena 422 (2021). doi: 10.1016/j.physd.2021.132902.

[59] J. Cox, R. Durrett, Limit theorems for the spread of epidemics and forest fires, Stochastic Processes and their Applications 30 (2) (1988) 171–191. doi:10.1016/0304-4149(88)90083-X.

[60] M. Lewis, S. Petrovskii, J. Potts, The Mathematics Behind Biological Invasions, Vol. 44, Springer International Publishing, Cham, 2016. doi: 10.1007/978-3-319-32043-4.

[61] C.-C. Chen, L.-C. Hung, M. Mimura, D. Ueyama, Exact travelling wave solutions of three-species competition–diffusion systems, Discrete and Continuous Dynamical Systems - B 17 (8) (2012) 2653–2669. doi:10.3934/dcdsb.2012.17.2653.

[62] J. Medlock, M. Kot, Spreading disease: integro-differential equations old and new, Mathematical Biosciences 184 (2) (2003) 201–222. doi: 10.1016/S0025-5564(03)00041-5.

[63] S. N. Busenberg, C. C. Travis, Epidemic models with spatial spread due to population migration, Journal of Mathematical Biology 16 (2) (1983) 181–198. doi:10.1007/BF00276056.

[64] T. C. Reluga, J. Medlock, A. P. Galvani, A Model of Spatial Epidemic Spread When Individuals Move Within Overlapping Home Ranges, Bulletin of Mathematical Biology 68 (2) (2006) 401–416. doi:10.1007/s11538-005-9027-y.

[65] M. Kot, M. A. Lewis, P. van den Driessche, Dispersal data and the spread of invading organisms, Ecology 77 (7) (1996) 2027–2042. doi: 10.2307/2265698.

[66] J. G. Skellam, Random dispersal in theoretical populations, Biometrika 38 (1-2) (1951) 196–218. doi:10.1093/biomet/38.1-2.196.

[67] M. Turelli, A. A. Hoffmann, Rapid spread of an inherited incompatibility factor in California Drosophila, Nature 353 (6343) (1991) 440–442. doi: 10.1038/353440a0.

[68] M. J. Tildesley, P. R. Bessell, M. J. Keeling, M. E. J. Woolhouse, The role of pre-emptive culling in the control of foot-and-mouth disease, Proceedings of the Royal Society B: Biological Sciences 276 (1671) (2009) 3239–3248. doi:10.1098/rspb.2009.0427.

[69] P. Rohani, X. Zhong, A. A. King, Contact network structure explains the changing epidemiology of pertussis., Science 330 (6006) (2010) 982–985. doi:10.1126/science.1194134.

[70] J. Mossong, N. Hens, M. Jit, P. Beutels, K. Auranen, R. Mikolajczyk, M. Massari, S. Salmaso, G. S. Tomba, J. Wallinga, J. Heijne, M. Sadkowska-Todys, M. Rosinska, W. J. Edmunds, Social Contacts and Mixing Patterns Relevant to the Spread of Infectious Diseases, PLOS Medicine 5 (3) (03 2008). doi:10.1371/journal.pmed.0050074.

[71] E. Volz, L. A. Meyers, Susceptible–infected–recovered epidemics in dynamic contact networks, Proceedings of the Royal Society B: Biological Sciences 274 (1628) (2007) 2925–2934. doi:10.1098/rspb.2007.1159.

[72] J. N. S. Eisenberg, M. A. Desai, K. Levy, S. J. Bates, S. Liang, K. Naumoff, J. C. Scott, Environmental determinants of infectious disease: a framework for tracking causal links and guiding public health research, Environmental health perspectives 115 (8) (2007) 1216–1223. doi:10.1289/ehp.9806.

[73] J. O. Lloyd-Smith, S. J. Schreiber, P. E. Kopp, W. M. Getz, Super-spreading and the effect of individual variation on disease emergence, Nature 438 (7066) (2005) 355–359. doi:10.1038/nature04153.

[74] J. D. Murray, E. A. Stanley, D. L. Brown, On the spatial spread of rabies among foxes, Proceedings of the Royal Society of London. Series B. Biological Sciences 229 (1255) (1986) 111–150. doi:10.1098/rspb.1986.0078.

[75] J. Paulose, J. Hermisson, O. Hallatschek, Spatial soft sweeps: Patterns of adaptation in populations with long-range dispersal, PLOS Genetics 15 (2) (2019) 1–29. doi:10.1371/journal.pgen.1007936.

[76] S. A. Lee, C. I. Jarvis, W. J. Edmunds, T. Economou, R. Lowe, Spatial connectivity in mosquito-borne disease models: a systematic review of methods and assumptions, Journal of The Royal Society Interface 18 (178) (2021) 20210096. doi:10.1098/rsif.2021.0096.

[77] Cdc.gov: How measles spreads [online] (11 2022) [cited 2024-11-13].

[78] D. M. Feehan, A. S. Mahmud, Quantifying population contact patterns in the United States during the COVID-19 pandemic, Nature Communications 12 (1) (2021) 893. doi:10.1038/s41467-021-20990-2.

[79] Census.gov: Urban areas facts [online] (10 2021) [cited 2023-06-04].

[80] Grenfell, B. T. and Bjørnstad, O. N. and Kappey, J., Travelling waves and spatial hierarchies in measles epidemics, Nature 414 (6865) (2001) 716–723. doi:10.1038/414716a.

[81] R. Durrett, S. Levin, The importance of being discrete (and spatial), Theoretical Population Biology 46 (3) (1994) 363–394. doi:10.1006/tpbi.1994.1032.

